# IPMK-1 (inositol phosphate multikinase) is required for optimal integrin adhesion complex assembly in *C. elegans* muscle

**DOI:** 10.1101/2025.09.09.675199

**Authors:** Sara Sagadiev, Robert Hudson, Yuchu Wang, Jordan Christmon, April Reedy, Hiroshi Qadota, Guy M. Benian

**Affiliations:** Department of Pathology, Emory University, Atlanta, Georgia 30322 USA

**Keywords:** muscle, integrin adhesion complexes, inositol phosphate multikinase, PIX pathway, genetic enhancer, C. elegans

## Abstract

Integrin adhesion complexes (IACs) are a network of many proteins including the transmembrane protein integrin that anchor cells to the extracellular matrix (ECM). In *C. elegans* muscle IACs exist at the bases of dense bodies and M-lines, and at the boundaries of adjacent muscle cells (MCBs). We have reported that the proper assembly of IACs requires the Rac GEF PIX-1 and members of the PIX-1 pathway. Here we report that further investigation into the original Million Mutation Project strain that led us to uncover the role of PIX-1 in muscle, revealed a genetic enhancer of the PIX-1 phenotype. Genetic mapping shows that the enhancer is a recessive mutation in a single gene, *ipmk-1*. *ipmk-1*, encodes inositol phosphate multikinase and converts PIP2 to PIP3, IP3 to IP4, and IP4 to IP5. Loss of function of *pix-1* results in reduced accumulation of IAC proteins at the MCB, and *ipmk-1; pix-1* double mutants additionally display a large gap between adjacent muscle cells. *ipmk-1* similarly enhances other members of the PIX pathway including PAK-1, and the RacGAP RRC-1. Lack of *ipmk-1* itself results in abnormal clumping of IAC components at the MCB, decreased whole animal locomotion, and mild sarcomere disorganization. Analysis of *ipmk-1; ipp-5* double mutants suggest that the MCB defect of *ipmk-1* is due to decreased PIP3 in the cell membrane rather than the decreased IP3 and Ca2+ signaling. Overall, these studies provide in vivo evidence, specifically in muscle, that the level of PIP3 and PIP2 influence the proper assembly of IACs.

**Article Summary:** *C. elegans* muscle have integrin adhesion complexes (IACs) at dense bodies, M-lines, and boundaries between adjacent muscle cells (MCBs). The Benian lab reported that assembly of these IACs requires the PIX-1 pathway. Here they report *ipmk-1* is a genetic enhancer of the PIX pathway. *ipmk-1*encodes inositol phosphate multikinase converting PIP2 to PIP3. *pix-1* shows reduced accumulation of IAC components; *ipmk-1; pix-1* double mutants additionally have large gaps between muscle cells. *ipmk-1* mutants display abnormal clumping of IAC components. This is the first report of the importance of these lipids in assembly of IACs in vivo, and in particular, in muscle.

## Introduction

Integrin adhesion complexes (IACs; aka focal adhesions), are assemblages of 50-100 different proteins including the transmembrane protein integrin which link the extracellular matrix (ECM) to the cytoskeleton and serve to anchor cells to the ECM (Bachir et al. 2014; Horton et al. 2015). In striated muscle, IACs are called costameres and link the peripherally located myofibrils to the muscle cell membrane and ECM and transmit the force of muscle contraction to the outside of the muscle cell (Ervasti, 2003; Henderson et al. 2017). The study of *C. elegans* continues to permit discovery of new aspects of muscle assembly, maintenance, regulation and aging (Gieseler et al. 2017). The major muscle in *C. elegans* is in the body wall and its cyclical contraction/relaxation permits movement of the whole animal (Benian and Epstein, 2011). In this muscle, IACs are found at 3 locations: the bases of the sarcomeric M-lines and dense bodies (Z-disks), and at the muscle cell boundaries (MCBs). MCBs consist of cell to ECM to cell and their IACs contain a subset of proteins located at dense bodies (Qadota et al. 2017). We reported (Moody et al. 2020), that the PIX signaling pathway is required for assembly of IACs at MCBs, by identifying and analyzing *pix-1* loss of function mutants. Moreover, despite having normally organized sarcomeres, *pix-1* null mutants display a ∼50% reduction in whole animal locomotion likely due to decreased transmission of lateral forces between adjacent muscle cells (Moody et al. 2020).

In mammals, the PIX pathway is important for development of nervous and immune systems (Schmalzigaug et al. 2009; Huang et al. 2011; Ramakers et al. 2012; Volinsky et al. 2006; Missy et al. 2008). In *C. elegans*, PIX-1 functions in the shape and migration of distal tip cells (Lucanic and Cheng, 2008), neuroblasts (Dyer et al. 2010) and tension-dependent morphogenesis of epidermal cells (Zhang et al. 2011). However, our lab was the first to show that the PIX pathway has a crucial function in muscle of any animal.

PIXs act as a Guanine Nucleotide Exchange Factors (GEFs) to promote exchange of GDP with GTP thereby activating small GTPases. In other organisms, PIXs activate Rac and Cdc42. We reported that *pix-1* mutants have reduced levels of GTP-bound(active) Rac in muscle (Moody et al. 2020). Activated (GTP bound) Rac binds to and activates PAK family protein kinases. Rho family GTPases (Rho, Rac, Cdc42) cycle between active (GTP bound) and inactive (GDP bound) states. In the PIX pathway, activation occurs by PIX. Inactivation is likely to occur via so called GTPase-activating proteins (GAPs), which promote the hydrolysis of GTP to GDP. In 2024, we reported identification of the first RacGAP for the PIX pathway in any organism or cell type, called RRC-1 (Moody et al., 2024). Loss of function of *rrc-1* results in reduced localization of IACs to the MCB, disorganization of M-lines and dense bodies, disorganization of sarcomeres, and reduced whole animal locomotion. Multiple lines of genetic and cell biologic evidence suggest that RRC-1 is part of the PIX-1 pathway: (1) each single mutant (*pix-1* or *rrc-1*) results in a similar defect at the MCB; (2) PIX-1 and RRC-1 each localize to the MCB; (3) localization of RRC-1 to the MCB depends on PIX-1 and vice versa; and (4) RRC-1 exists in a complex with PIX-1.

Here we report further studies of the original Million Mutation Project (Thompson et al. 2013) strain that led us to identify *pix-1* (Moody et al. 2020). We found that the original strain has a genetic enhancer of the PIX phenotype, and that the enhancement results from loss of function of a single recessive gene. The enhancer gene is *ipmk-1*, which encodes inositol phosphate multikinase, a highly conserved enzyme that converts the phosphoinositides PIP2 to PIP3, and the inositol polyphosphates IP3 to IP4, and IP4 to IP5. The IPs are involved in calcium signaling, whereas the PIPs are incorporated into cell membranes. Loss of function of *ipmk-1* itself results in an abnormal clumping of IAC components at the MCB. Based on genetic evidence we strongly suspect that the muscle phenotype of *ipmk-1* is due to either reduced levels of PIP3 or increased levels of PIP2 in the muscle cell membrane, rather than through IP3 and Ca^2+^ signaling. Overall, we provide genetic evidence that the metabolism of phosphoinositides in muscle influences the proper assembly of IACs, possibly through interaction with the transmembrane domains of integrins and PIP-binding domains of several key IAC proteins.

## Materials and Methods

### *C. elegans* strains

All but one nematode strain was grown on NGM plates using standard methods and maintained at 20°C (Brenner, 1974). The exception was *ipmk-1(tm2687)*, which for some immunostaining was grown on NGM plates containing 50 μM 5-fluoro-2’-deoxyuridine (FUDR). This was done because *ipmk-1(tm2687)* grows slowly, and although for most strains we image muscle at day 1 or 2 of adulthood, at that age *ipmk-1(tm2687)* adults are small and consequently have small body wall muscle cells. If we wait until day 4 or 5, when the muscle cells are bigger, then embryos push up against the body wall and distort our images. By including the DNA replication inhibitor FUDR adults do not develop embryos, and this imaging problem is avoided. Most strains were obtained from the Caenorhabditis Genetics Center. The wild type strain was N2 (Bristol).

VC20386, contains *pix-1(gk299374)* and *ipmk-1(gk214790)*

GB291, *pix-1(gk299374)* derived from VC20386 and outcrossed 5X to wild type GB398, “chromosome III GFP”, *sfIs37* [hsp::HA::CUL-1; sur-5::GFP]

GB371, “chromosome IV GFP”, *sfIs27* [hsp::UNC-45::HA; sur-5::GFP] *ipmk-1(tm2687)*

GB397, *ipmk-1(tm2687)* outcrossed 6X to wild type

GB399, VC20386 outcrossed 4X to *pix-1(gk299374)*, 5X OC, resulting in maintenance of 11 of 13 mutations on IV (as compared to wild type); note: other chromosomes are wild type except for the *pix-1* mutation on X

GB400, GB399 harboring sfEx81 [myo-3p::HA::IPMK-1; sur-5::GFP]

The following strains were crossed into GB397 [*ipmk-1(tm2687* 6X OC)] to make double mutants:

- RB689, pak-1(ok448)
- GB340, *pak-1(syb647)* outcrossed 4X to wild type
- GB343, *rrc-1(ok1747)* outcrossed 5X to wild type
- PS3653, *ipp-5(sy605)* outcrossed 7X to wild type

The following strains were crossed into GB291 [*pix-1(gk299374)*], to make double mutants:

- *alp-1(tm1137)*
- RB2198, C27B7.7(ok2978)
- VC2141, C27B7.7(ok2868)
- VC40162 contains K08E7.6(gk491604)
- F07F5.8(tm6279)
- F07F5.8(tm6114)

### Immunostaining and confocal microscopy of body wall muscle

Adult worms were fixed and immunostained following the protocol of Nonet et al. (1993), with details described by Wilson et al. (2012). Unless otherwise stated, most primary antibodies were used at a 1:200 dilution. These included: anti-PAT-6 (rat polyclonal; used at 1:100 dilution; Warner et al., 2013), anti-UNC-52 (mouse monoclonal MH2; Mullen et al., 1999; obtained from the University of Iowa Hybridoma Bank), anti-UNC-95 (rabbit polyclonal Benian-13; used at 1:100 dilution; Qadota et al., 2007), anti-UNC-112 (used at 1:100 dilution; Hikita et al., 2005), anti-MHC A (mouse monoclonal 5–6; Miller et al., 1983; from the University of Iowa Hybridoma Bank), anti-UNC-89 (mouse monoclonal MH42; Hresko et al. 1994; Benian et al., 1996), and anti-ATN-1 (mouse monoclonal MH35; Francis and Waterston, 1991; generously gifted by Pamela Hoppe, Western Michigan University). Secondary antibodies (anti-rabbit Alexa 488, anti-rat Alexa 594, and anti-mouse Alexa 594; Invitrogen) were also used at 1:200 dilution. Fixation and staining with phalloidin-rhodamine were performed according to Waterston et al. (1984). Imaging was performed at room temperature using a Zeiss LSM510 confocal system with an Axiovert 100M microscope and a 63×/1.4 NA oil immersion Apochromat objective in 1× mode, except for Figure 12C. The images shown in Figure 12C were taken with the same objective but in the 2.5x zoom mode. All confocal images were processed for color balance using Adobe Photoshop (Adobe, San Jose, CA).

### Transgenic rescue of enhanced (gap) phenotype of ChrIV(mut); *pix-1* strain by muscle expression of wild type IPMK-1

To create pPD95.86-HA-IPMK-1, a full-length cDNA for wild type IPMK-1 was generated by using PCR with the following primers and the RB2 cDNA library: ipmk-1_rescue-F:

GCGGTTAACATGAGTGTGCTATCAAATGAC (begins with added HpaI-ATG) ipmk-1_rescue-R:

CGCGTCGACCTATGAAGAAATAATATTATTC (ends with TAG stop and SalI)

The amplified fragment was cloned into the EcoRV and SalI sites of pKS-HA(Nhex2) and the sequence was determined to be error-free by Sanger sequencing, resulting pKS-HA-IPMK-1. The NheI fragment of pKS-HA-IPMK-1 was cloned into pPD95.86 NheI site, resulting pPD95.86-HA-IPMK-1, expressing HA tagged IPMK-1 under the control of the *myo-3* promoter.

To create GB400, pPD95.86-HA-IPMK-1 was injected into 4x outcrossed VC20386 (GB399, ChrIV(mut); *pix-1(gk299374*)) strain with sur-5-gfp (pTG96) as the transformation marker (Yochem et al. 1998), and screening for GFP positive worms and establishing a transgenic line.

### Swimming and crawling assays

Swimming and crawling assays on Day 1 or Day 2 adults were carried out as described in Moody et al. (2024).

### Generation of antibodies to IPMK-1 and western blots

Rabbit polyclonal antibodies were generated to the N-terminal 120 residues of IPMK-1. The coding sequence of this region was generated by PCR from a cDNA library (RB2), and cloned into vectors pGEX-KK1 and pMAL-KK1, to express glutathione-S-transferase (GST) and MBP fusion proteins in E. coli strain Rosetta 2 (DE3)(Millipore Corporation, cat. no. 71397-4). Protein production and purification was carried essentially as described in Matsunaga et al. (2024). GST-IPMK-1 (1-120) was shipped to Pocono Rabbit Farm and Laboratory Inc. (Canadensis, PA) as the antigen for generation of polyclonals in 2 rabbits. Antibodies were affinity-purified essentially as described previously (Matsunaga et al. 2024) using the MBP-IPMK-1 (1-120) antigen, but because the pI of this fusion protein is 5.22, it was coupled to Affigel-15 (Bio-Rad Laboratories, cat. no. 153-6052) which is best suited for coupling acidic proteins. A western blot was used to test the specificity of the anti-IPMK-1 antibodies. The method of Hannak et al. (2002) was used to prepare total protein lysates from wild type, *ipmk-1(gk214790)* [from GB399] and *ipmk-1(tm2687).* Proteins were separated on a 10% SDS-PAGE gel, transferred to nitrocellulose and reacted against affinity-purified anti-IPMK-1 at 1:1500 dilution which had been pre-absorbed against an acetone powder of E. coli OP50 to reduce reaction to bacterial proteins, followed by ECL and detection by film. For the western blot shown in Figure 7A demonstrating transgenic expression of HA-IPMK-1 under the control of the *myo-3* promoter, an extract was prepared, separated by SDS-PAGE, transferred to nitrocellulose membrane as described above, and reacted against anti-HA (rabbit monoclonal C29F4; Cell Signaling Technology) at 1:1,000 dilution, and detected by ECL and film exposure.

## Results

The *pix-1* gene was discovered to be involved in the assembly or stability IAC components at the muscle cell boundary (MCB) by screening of the Million Mutation Project (MMP) collection of adult mutants (Moody et al., 2020). We reported that the MMP strain VC20386 shows an abnormality at the muscle cell boundary (MCB) but no defects at dense bodies and M-lines, the other two structures in muscle cells that are based on IACs. After outcrossing VC20386 3 times against wild type animals, we genetically mapped the MCB defect to the single gene, *pix-1*, and found similar defects in 4 additional independently-isolated *pix-1* mutants (Moody et al. 2020). However, we observe that the original VC20386 strain shows a more severe phenotype, than the strains that had been outcrossed to wild type, or other *pix-1* mutants, showing a large gap between adjacent body wall muscle cells (indicated by arrow in Figure 1), in addition to loss of accumulation of PAT-6 (Figure 1) and other IAC components.

**Figure 1.**
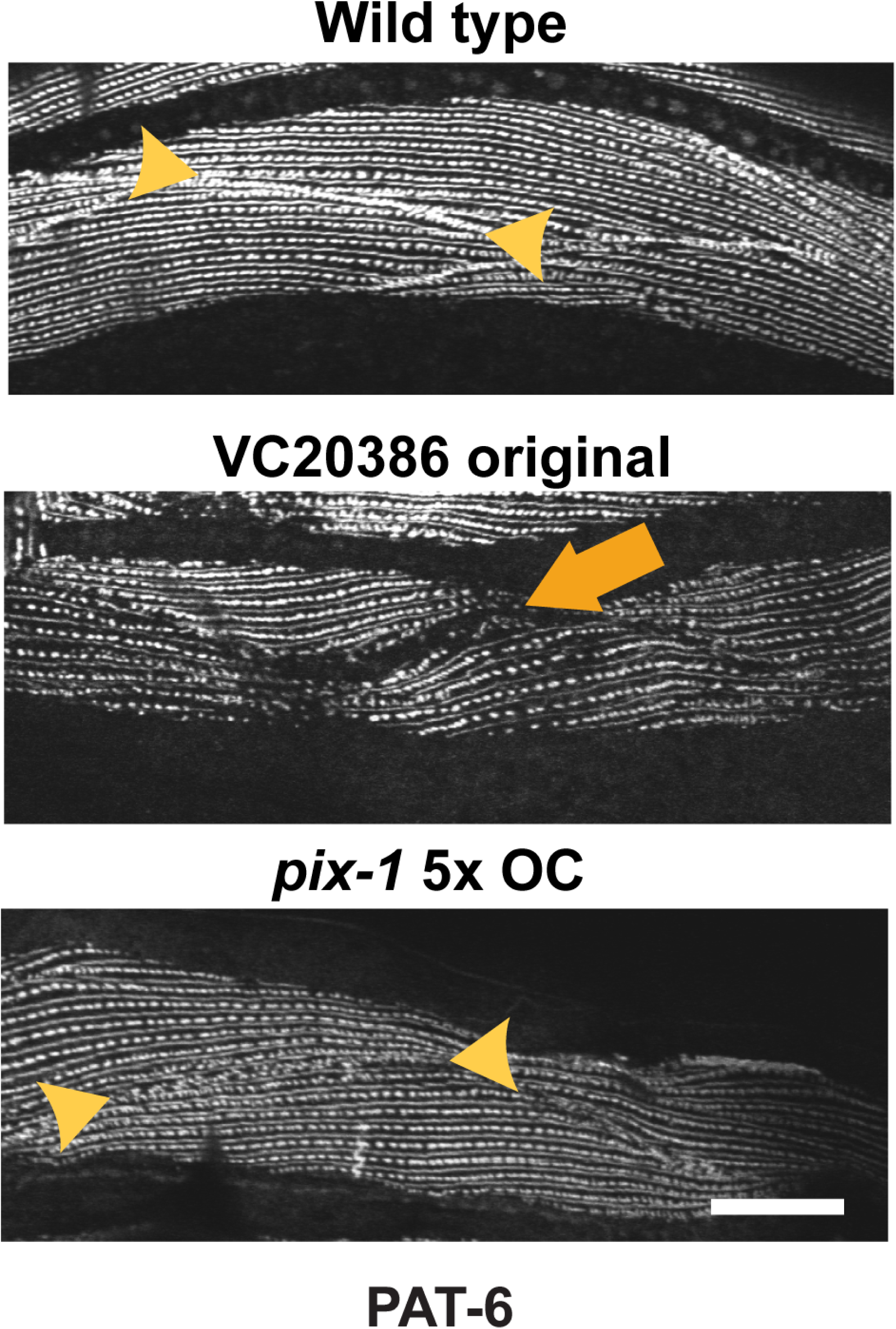
The MMP strain, VC20386, from which the *pix-1* mutant was isolated shows a more severe phenotype than *pix-1* itself. Immunostaining of wild type with anti-PAT-6 shows the accumulation of PAT-6 at the MCB (indicated by arrowheads). The *pix-1(gk299374)* mutant derived from VC20386 after 5 outcrosses to wild type shows lack of accumulation of PAT-6 at the MCB. However, the original VC20386 strain shows lack of accumulation at the MCB and a large gap between adjacent body wall muscle cells (indicated by arrow). Scale bar, 10 µm.

After outcrossing VC20386 just one time to wild type, this “big gap” phenotype was not seen, suggesting that the original VC20386 contains an additional extragenic enhancer mutation. To test this hypothesis, we crossed the original VC20386 strain with the 5x outcrossed *pix-1* strain (derived from VC20386). Actually, this cross was done with the 5X outcrossed *pix-1* strain having sur-5::gfp inserted in the left arm of Chromosome III, denoted as, GFP III; *pix-1* X 5x OC. The GFP was used as a marker for following the cross. First, we crossed GFP III; *pix-1* X 5x OC with wild type males (+/+ III; +/- X), and then picked up GFP males (GFP/+ III; *pix-1*/- X), and these GFP males were crossed to the original VC20386 (Figure 2A). We picked up GFP F1 hermaphrodites (Figure 2b), allowed them to self-fertilize, and selected 12 non-GFP F2 worms. In all these F2 animals, the *pix-1* mutation was maintained (Figure 2c). We collected and fixed several hundred F3 progeny from these 12 F2 worms, and immunostained them with anti-PAT-6 (Figure 3). Among the 12 F2s, 2 F2 animals displayed the “big gap” phenotype (indicated by large orange arrows in Figure 3), suggesting the presence of an additional mutation that is a recessive enhancer, “en”, of *pix-1*.

**Figure 2.**
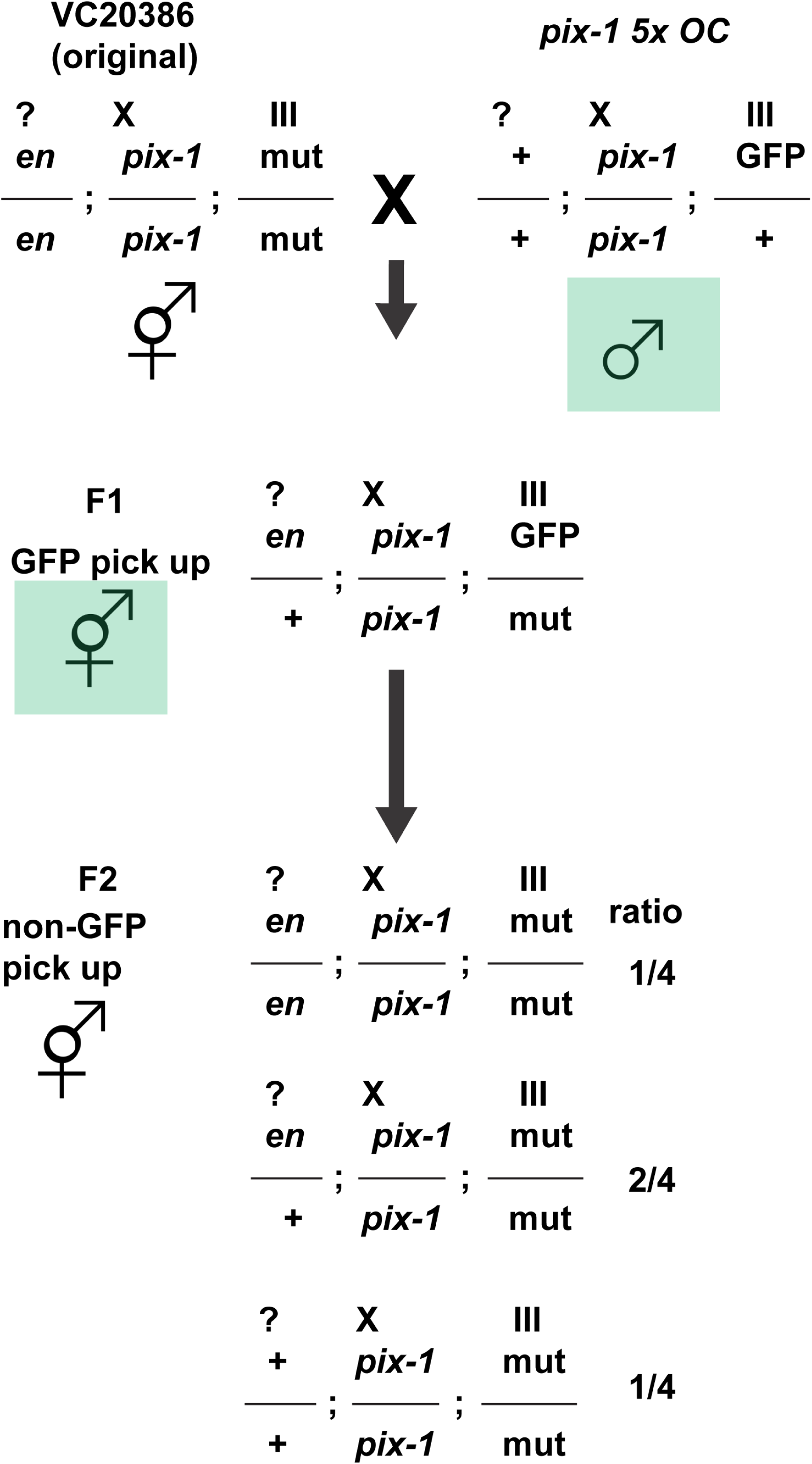
Crossing scheme used to test the hypothesis that the original strain VC20386 has a recessive mutation in an additional gene that enhances the phenotype of *pix-1*.

**Figure 3.**
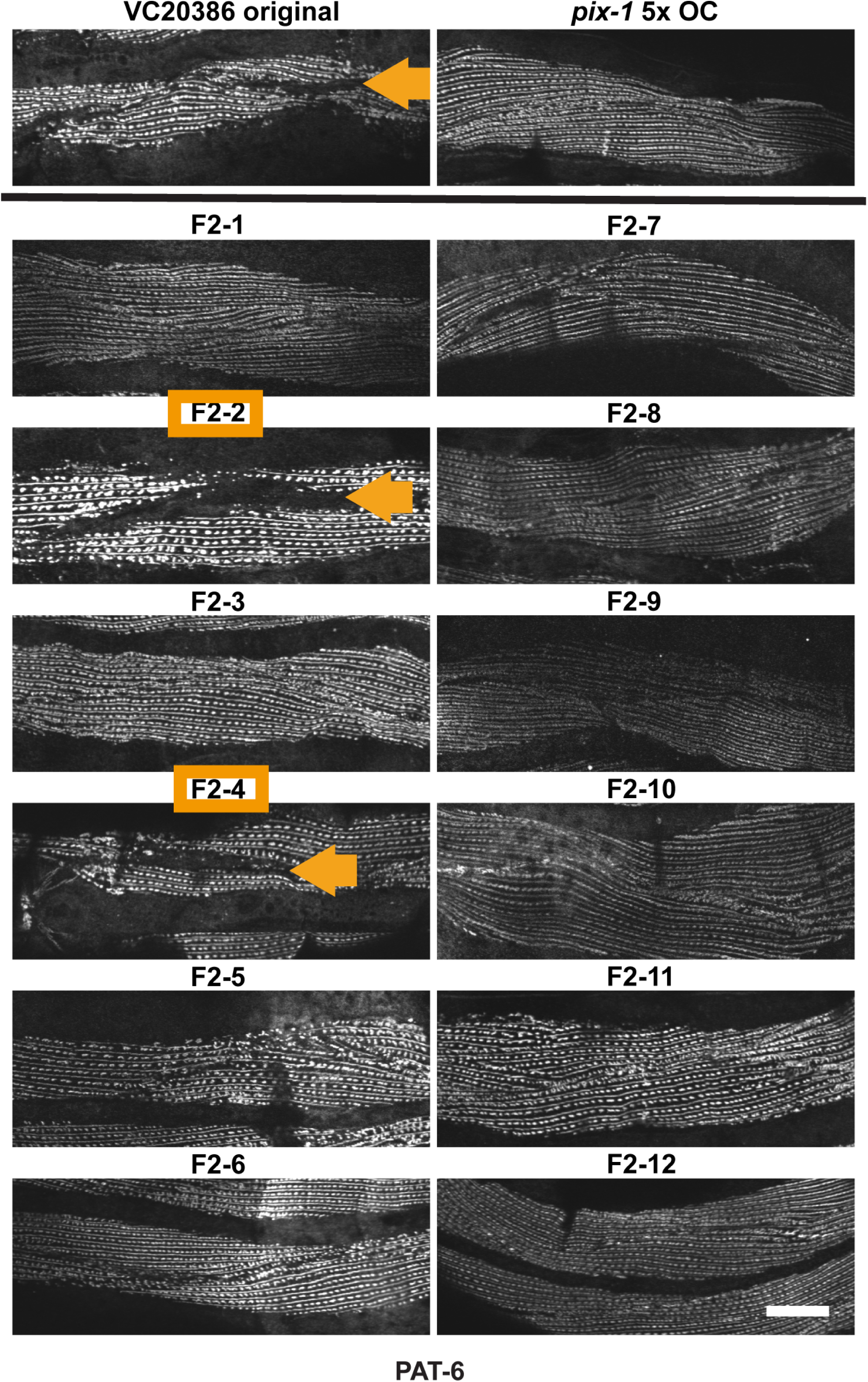
The enhancer mutation resides in a single gene and is recessive. Images of anti-PAT-6 immunostaining of body wall muscle with anti-PAT-6. On top, is shown the original VC20386 strain and the *pix-1* mutant derived from it after 5 outcrosses to wild type. Below are shown immunostaining of 12 randomly selected F2 animals from the crosses depicted in Figure 2. Note that the large gap or enhanced phenotype (denoted by arrows) is found in 2 out of 12 animals, which is close to the expected ratio of ¼ if the enhancer mutation is recessive. Scale bar, 10 µm.

One of these 2 F2 animals with the big gap, en/en; *pix-1/pix-1* X, was crossed to GFP III; *pix-1* X 5x OC two more times, to recover en/en; *pix-1/pix-1* X 3x OC. (“OC” meaning outcrossed to wild type). Since the MMP has revealed all the mutation sites of VC20386 (Thompson et al. 2013), we selected 12 mutations as SNP markers, one for the right and left end of each of the 6 nematode chromosomes (Supplementary Table 1), to allow us to distinguish by PCR and DNA sequencing, the original VC20386 strain from wild type. After sequencing the 12 SNP markers from en/en; *pix-1/pix-1* X 3x OC, we determined that the left arm of chromosome III and left arm of chromosome IV remained mutant (Figure 4).

**Figure 4.**
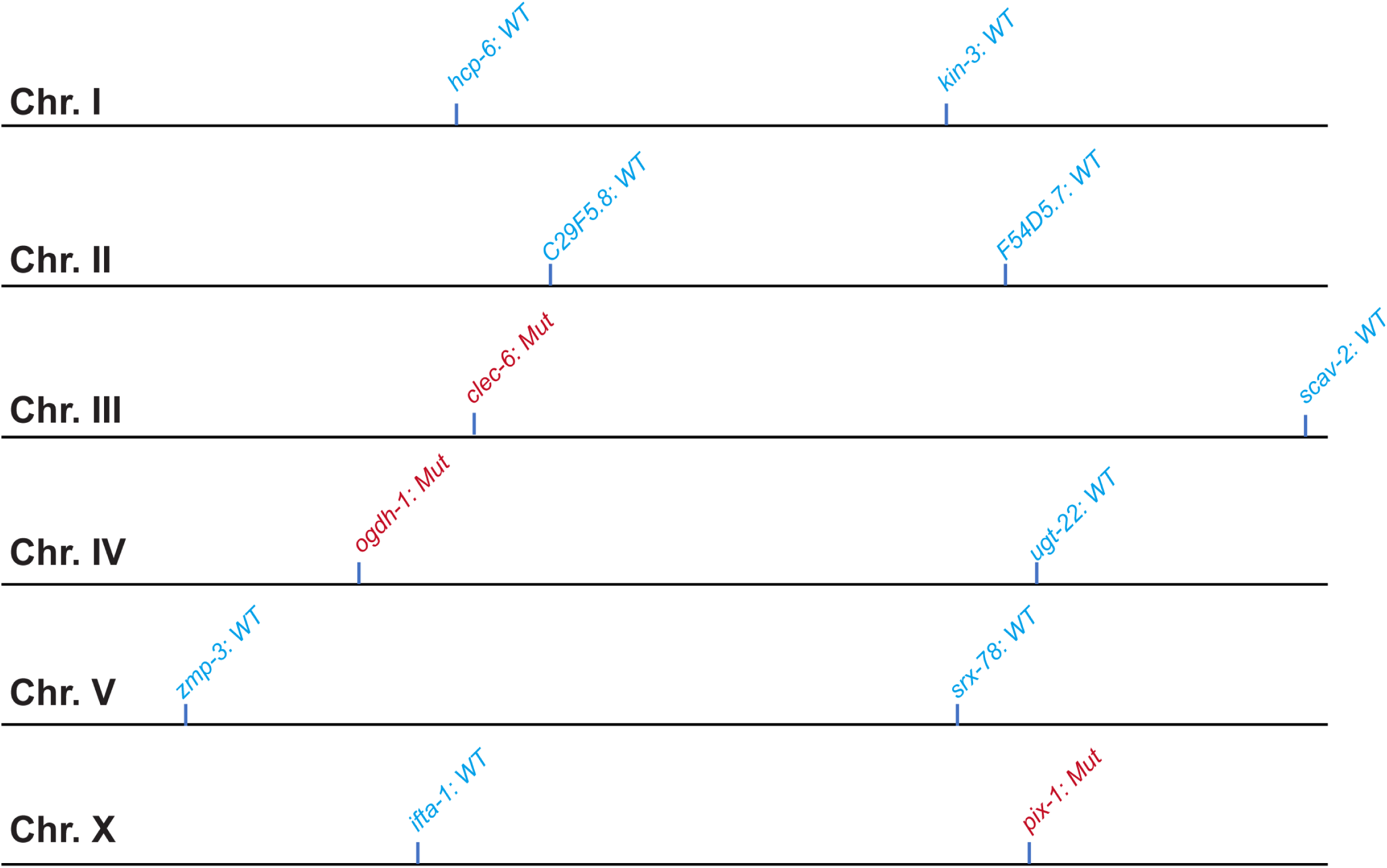
Initial mapping of the enhancer mutation to chromosomes III or IV. One of the 2 F2 animals with the big gap, en/en; *pix-1/pix-1* X, was crossed to GFP III; pix-1/pix-1 X 5x OC two more times, to recover en/en; *pix-1/pix-1* X 3x C. Since the MMP has revealed all the mutation sites of VC20386 (Thompson et al. 2013), we selected 12 mutations as SNP markers, one for the right and left end of each of the 6 nematode chromosomes, to allow us to distinguish by PCR and DNA sequencing, the original VC20386 strain from wild type (SNP from wild type in blue, SNP from VC20386 in red). After sequencing the 12 SNP markers from en/en; *pix-1/pix-1* X 3xC, we determined that the left arm of chromosome III and left arm of chromosome IV remained mutant (indicated by red font).

Since we used a GFP insertion in chromosome III to follow our crosses, we next crossed en/en; *pix-1/pix-1* X 3x OC with a strain in which GFP had been inserted in chromosome IV, and thus eliminating mutations on chromosome III. Upon immunostaining, the resulting strain still showed the “big gap” phenotype (Supplemental Figure 1), suggesting that the enhancer mutation does not reside on chromosome III but that the enhancer mutation resides on the left arm of chromosome IV.

The original VC20386 strain has exonic SNPs in 13 genes on chromosome IV (from *col-101* through Y73F8A.33). When we sequenced *ogdh-1* and *ugt-22* from the en/en; *pix-1/pix-1* 3XOC strain, we found that *ogdh-1* remained mutant, but *ugt-22* became wild type (Figure 4 and top of Figure 5). To map the enhancer mutation more finely, we sequenced 3 more of the SNPs (Supplementary Table 2) in this region and found that SNPs in the *sdz-27*, *wdr-46* and K08E7.6 genes were mutant. At this point, we concluded that the enhancer mutation was located to the left of *ugt-22* on IV, precisely the nucleotide 1-12580365 region (second line, Figure 4). This region of VC20386 contains 11 exonic SNPs in 11 genes (Supplementary Table 3). Within this region we found that, based on SAGE data (Meissner et al., 2009), 4 genes are expressed in muscle (*alp-1*, C27B7.7, *ipmk-1,* and K08E7.6), and one gene’s SNP is a nonsense mutation (K07F5.8)(last line of Figure 5).

**Figure 5.**
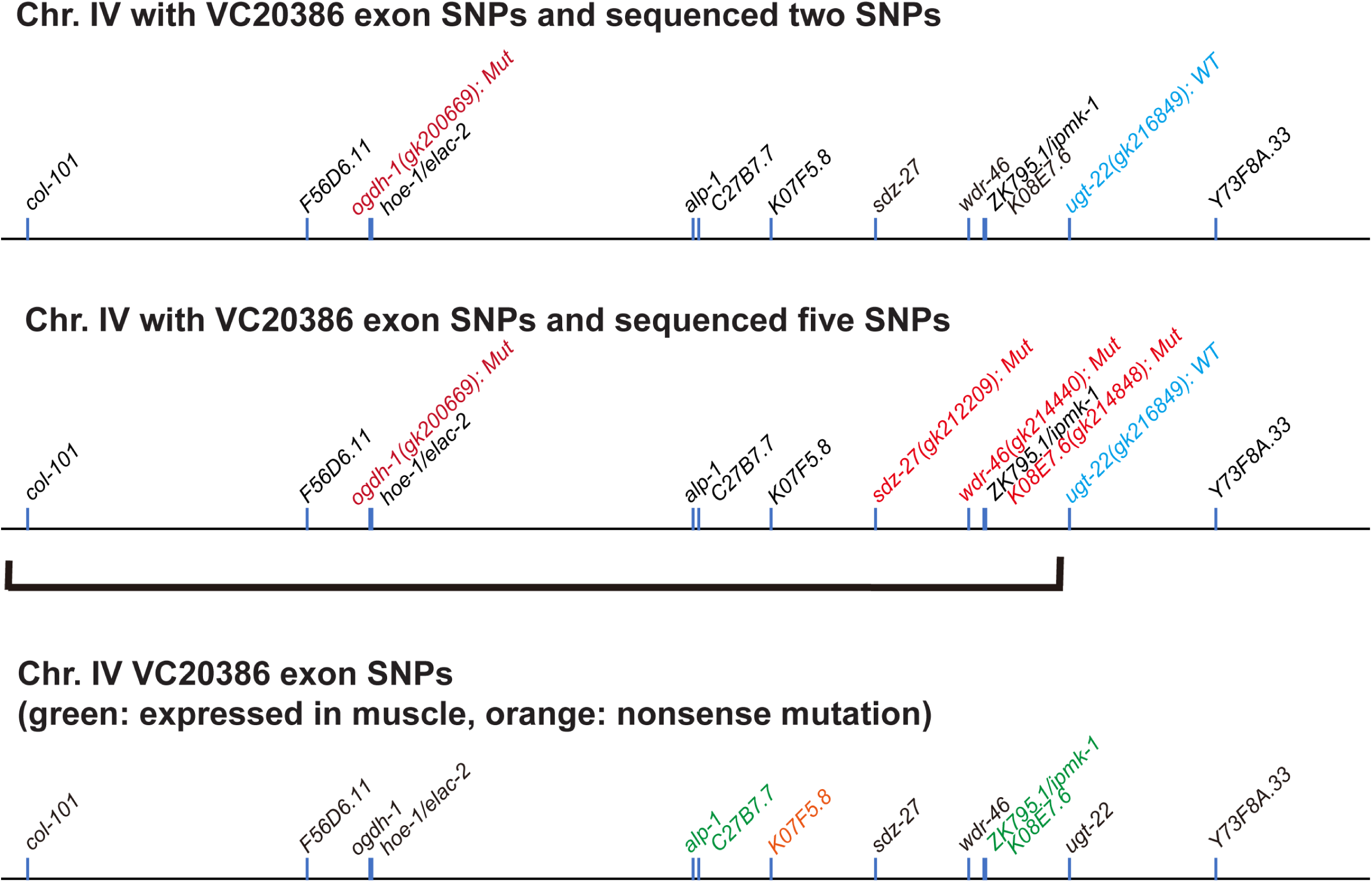
Mapping the enhancer to 5 candidate genes on chromosome IV. The original VC20386 strain has exonic SNPs in 13 genes on chromosome IV (from *col-101* through Y73F8A.33). When we sequenced *ogdh-1* and *ugt-22* from the en/en; pix-1/pix-1 3XOC strain, we found that *ogdh-1* remained mutant, but *ugt-22* became wild type (top map). To map the enhancer mutation more finely, we sequenced 3 more of the SNPs in this region and found that SNPs in the *sdz-27*, *wdr-46* and K08E7.6 genes were mutant. We concluded that the enhancer mutation was located to the left of *ugt-22* on IV, precisely the nucleotide 1-12580365 region (middle map). This region of VC20386 contains 11 exonic SNPs in 11 genes. Within this region we found that, based on SAGE data (Meissner et al., 2009), 4 genes are expressed in muscle (*alp-1*, C27B7.7, *ipmk-1*, and K08E7.6), and one gene’s SNP is a nonsense mutation (K07F5.8)(bottom map). Blue font: wild type sequence; red font: mutant (VC20386) sequence. Green font: gene is expressed in body wall muscle; orange font: gene with nonsense mutation in VC20386.

To determine which of these 5 genes is the enhancer gene, we obtained independently-generated mutations for each and made double mutants with *pix-1 (gk299374)* 5X OC, and performed immunostaining with anti-PAT-6 antibodies. As shown in Figure 6A, the double mutant *ipmk-1(tm2687); pix-1(gk299374)*, displays the big gap phenotype, like the VC20386 original strain. In addition, double mutants with the other 4 genes failed to show the big gap phenotype (Supplementary Figure 2). As shown in Figure 6B, the gap is also found when the double mutant is immunostained with antibodies to 2 additional IAC components, UNC-95 and UNC-52. As indicated in Figure 6C, the *ipmk-1* mutation in the original strain VC20386, called *gk214790*, is a GtoA mutation that disrupts a normal splicing site before the terminal coding exon, whereas *tm2687* is a 190 bp intragenic deletion that removes part of exon 3 and all of exon 4. Thus, both alleles of *ipmk-1*, *gk214790* and *tm2687,* are loss of function mutations.

**Figure 6.**
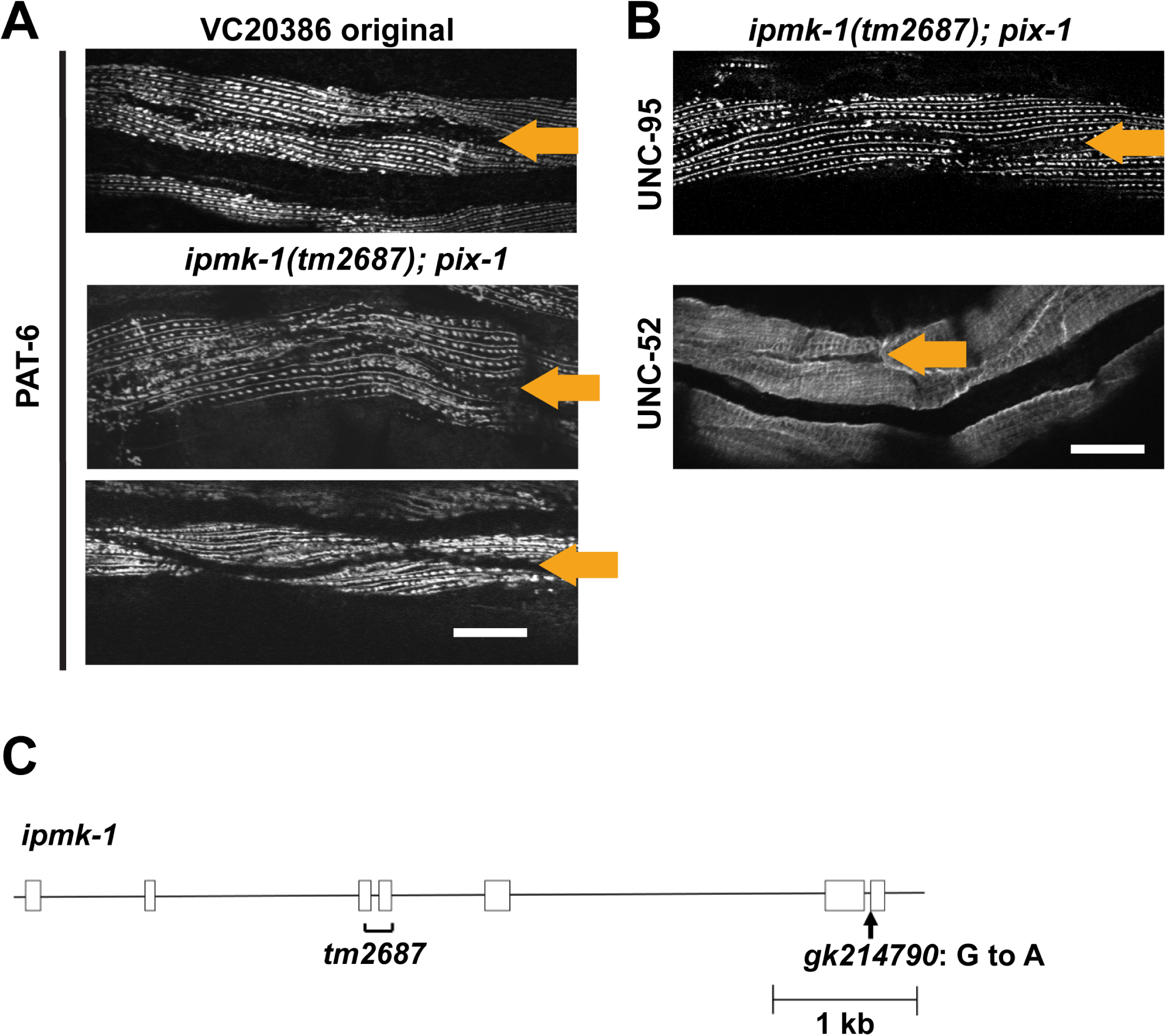
Double mutant analysis shows that the enhancer gene is *ipmk-1*. A. To determine which of the 5 genes is the enhancer gene, we obtained independently generated mutations for each and made double mutants with *pix-1(gk299374)* 5X OC, and performed immunostaining with anti-PAT-6 antibodies. The double mutant *ipmk-1(tm2687); pix-1(gk299374)*, displays the big gap phenotype, like the VC20386 original strain. Double mutants with the other 4 genes failed to show the big gap phenotype (Supplemental Figure 2). Two typical examples of body wall muscle from the double mutant are shown. B. The big gap phenotype of the double mutant is also found when immunostained with 2 additional IAC proteins, UNC-95 and UNC-52. C. Location of two *ipmk-1* mutant alleles. Displayed is an exon-intron map of the *ipmk-1* gene obtained from WormBase. *ipmk-1(tm2687)* is a 190 bp deletion that removes part of exon 3 and all of exon 4. *ipmk-1(gk214790)*, which exists in the original strain VC20386, is a G to A mutation that disrupts the normal splicing site before the terminal coding exon. Orange arrows indicate gaps between muscle cells. Scale bars, 10 µm.

To obtain additional evidence that *ipmk-1* is the enhancer mutation, we performed a transgenic rescue experiment. We created a plasmid designed to express an HA tagged wild type copy of IPMK-1 under the control of a nearly muscle-specific promoter for *myo-3* (myo-3p-HA-IPMK-1) and created a transgenic line in the en/en; *pix-1/pix-1* 4XOC strain, which contains mutations in 11 genes, including *ipmk-1*. This transgenic shows specific expression of HA-IPMK-1 by western blot (Figure 7A), and upon immunostaining shows rescue of the big gap phenotype (Figure 7B), that is, it resembles *pix-1* by itself. This is further evidence that the *pix-1* enhancer is *ipmk-1*.

**Figure 7.**
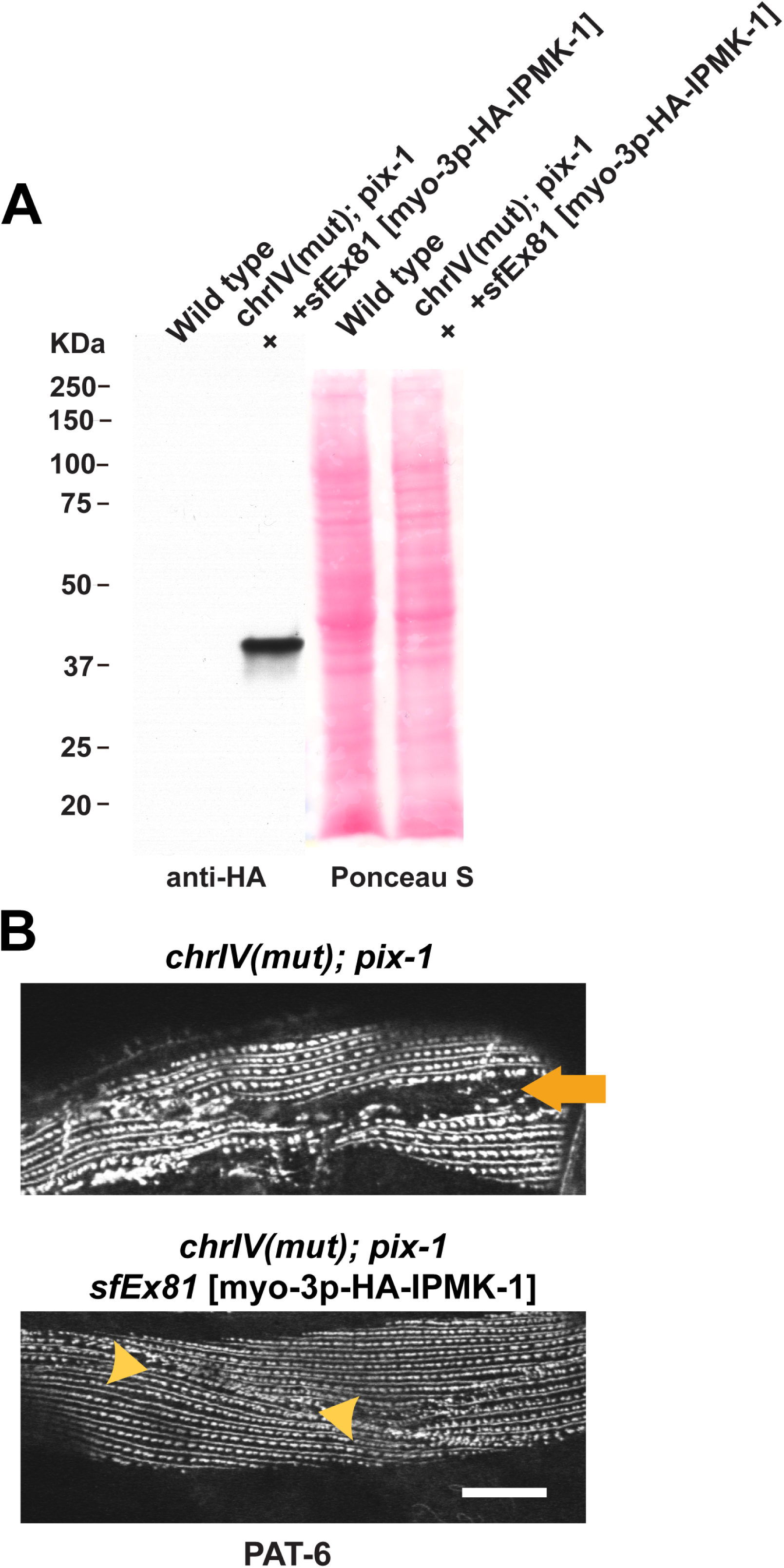
Muscle expression of *ipmk-1* rescues the big gap phenotype. As additional evidence that *ipmk-1* is the enhancer mutation, we performed a transgenic rescue experiment. We created a plasmid that would express an HA tagged wild type IPMK-1 under the control of a nearly muscle-specific promoter (myo-3p-HA-IPMK-1) and created a transgenic line in the en/en; pix-1/pix-1 4XOC strain (GB399), which contains mutations in 11 genes, including *ipmk-1.* This transgenic shows specific expression of HA-IPMK-1 by western blot (A), and upon immunostaining shows rescue of the big gap phenotype (B), that is, it resembles *pix-1* by itself. Orange arrow points to gap between muscle cells; arrow heads indicate MCB with lack of accumulation of PAT-6. Scale bar, 10 µm.

*ipmk-1* encodes the enzyme inositol phosphate multikinase (IPMK-1), conserved from yeast to humans (Kim et al. 2017). IPMK-1 converts PIP2 to PIP3, and IP3 to IP4, and IP4 to IP5. IP3 is involved in calcium signaling, whereas the PIPs are incorporated into cell membranes. Loss of function of *ipmk-1* has been reported to result in retarded postembryonic growth and a prolonged defecation cycle period in *C. elegans* and to do so through IP3/ Ca2+ signaling (Yang et al. 2021). However, there were no reports of *ipmk-1* having a function in muscle.

We next wondered if *ipmk-1* could enhance mutations in other members of the known PIX pathway. To examine this question, we prepared a double mutant with a loss of function mutation in a gene that encodes the known output of the Pix pathway, PAK-1, and with a loss of function mutation in a gene that encodes the RacGAP of the PIX pathway, RRC-1 (Moody et al. 2024). As shown in Figure 8, the *ipmk-1(tm2687); pak-1(ok448)* double, when immunostained with anti-PAT-6, displays the big gap phenotype like *ipmk-1; pix-1* mutants (Figures 6 and 7). Intriguingly, however, *ipmk-1(tm2687); rrc-1(ok1747)* shows both a big gap and assembly of IAC zipper-like structures (Figure 8). We have observed these zipper-like structures in wild type muscle (Qadota et al. 2017) only with super-resolution SIM which has twice the resolution of conventional confocal microscopy, and yet the image shown in Figure 8 was taken with confocal microscopy. Thus, the zipper-like IAC structures in the *ipmk-1; rrc-1* double mutant are bigger than those formed in wild type muscle. Finally, we also made an *ipmk-1(tm2687); pak-1(syb647)* double mutant in which *pak-1* has a L99F mutation which is predicted to make the PAK-1 protein kinase constitutively active (Moody et al. 2024). The *ipmk-1(tm2687); pak-1(syb647)* show an intermediate phenotype, with a smaller gap and smaller but still larger than wild type zipper-like structures. Therefore, loss of function of *ipmk-1* can enhance the phenotype of mutants in multiple proteins within the PIX pathway. Given that PIX pathway activity is reduced in *pix-1* and *pak-1(ok448)*(Moody et al. 2020), increased in *rrc-1(ok1747)* and *pak-1(syb647*)(Moody et al. 2024), this suggests that the formation of IACs at the MCB is promoted by increased activity of the PIX pathway.

**Figure 8.**
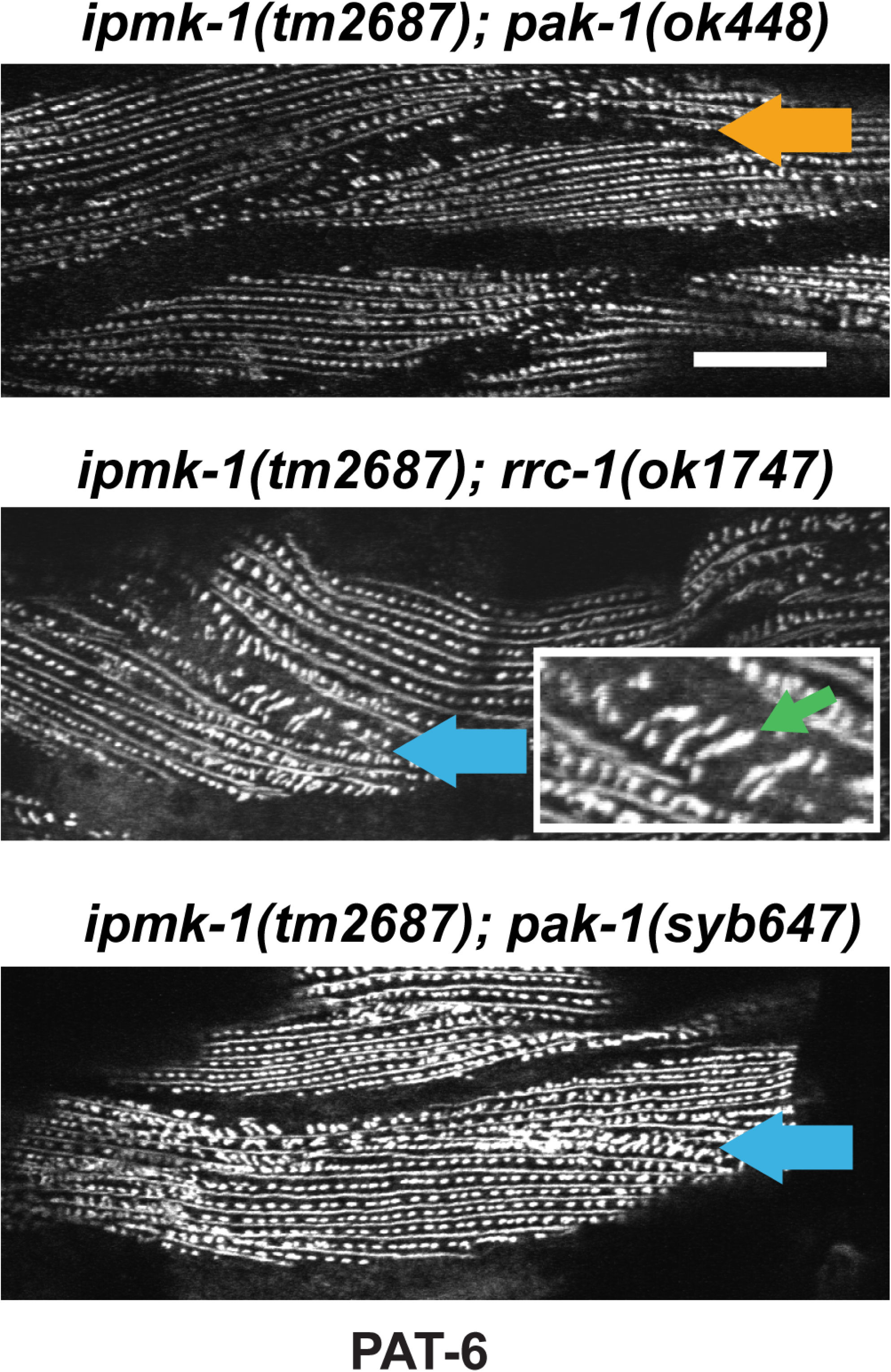
*ipmk-1* also enhances the muscle phenotype of *pak-1* and *rrc-1*, and MCB zipper-like structures are promoted by increased activity of the PIX pathway. The indicated double mutants were immunostained with anti-PAT-6. *ipmk-1; pak-1(ok448)* which includes a *pak-1* null mutant shows a phenotype very similar to *ipmk-1; pix-1*, with big gaps between muscle cells (orange arrow). However, *ipmk-1; rrc-1* which includes a *rrc-1* null mutant shows big gaps and assembly of IAC zipper like structures (blue arrow). The inset is a magnified view of part of the MCB, with a green arrow pointing to one of the zipper “teeth”. We have observed these zipper-like structures in wild type only with super-resolution SIM, and yet the image shown was taken with conventional confocal microscopy. Thus, the zipper-like structures in *ipmk-1; rrc-1* double mutants are bigger than in wild type. *ipmk-1; pak-1(syb647)* which includes a *pak-1* constitutively active kinase mutant shows an intermediate phenotype, with a smaller gap and smaller but still visible zipper-like structures. Scale bar, 10 µm.

*ipmk-1(tm2687)* is likely to be a null allele of *ipmk-1* since it is an intragenic deletion, removing 190 bp of sequence, including part of exon 3 and the entire exon 4 (Figure 6C). We outcrossed this strain to wild type 3 times and performed immunostaining with antibodies to IAC components. As shown in Figure 9, *ipmk-1(tm2687)* displays abnormal clumping or large aggregates of PAT-6, UNC-112 and UNC-95, but not UNC-52 at the MCB, both close to the outer muscle cell membrane, and even more so, deeper into the cell (indicated by red arrows in Figure 9). The fact that the ECM component UNC-52 (perlecan) is not affected by loss of IPMK-1 (i.e. UNC-52 shows uniform localization throughout the MCB), is further indication that the role of IPMK-1 is at the muscle cell membrane.

**Figure 9.**
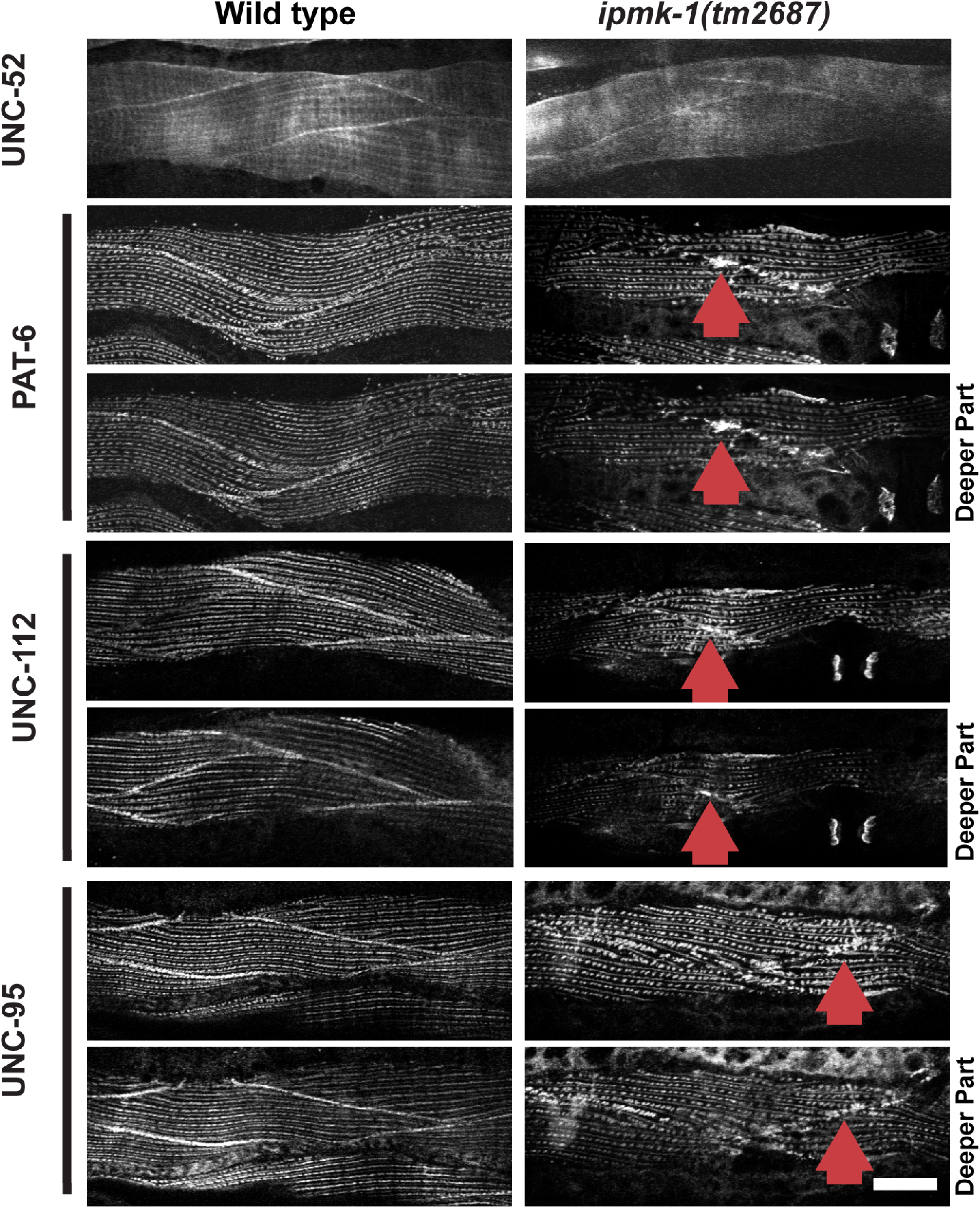
*ipmk-1(tm2687)* displays abnormal clumping of IAC components at the muscle cell boundary (MCB). Wild type and *ipmk-1(tm2687)* were immunostained with antibodies to UNC-52, PAT-6, UNC-112, and UNC-95, and imaged by confocal microscopy. In each case images were taken close to the outer muscle cell membrane and deeper into the muscle cell (“Deeper Part”). Note that in wild type these IAC proteins are uniformly distributed along the length of the MCB. However, *ipmk-1(tm2687)* shows abnormal clumping or aggregates of PAT-6, UNC-112 and UNC-95, but not UNC-52, at the MCB (indicated by red arrows). Scale bar, 10 µm.

Loss of function of two components of the PIX pathway affect sarcomere organization differently. Whereas loss of function of *pix-1* has normal sarcomere structure (Moody et al. 2020), loss of function of *rrc-1* results in moderately disorganized sarcomeres including I-bands, A-bands, M-lines and dense bodies (Moody et al. 2024). Thus, we wondered what effect *ipmk-1* would have on sarcomere organization. As shown in Figure 10, *ipmk-1(tm2687)* does not affect the organization of I-bands (phalloidin staining) or the main portions of dense bodies (ATN-1 staining) but does mildly affect the organization of A-bands (MHC A staining) and M-lines (UNC-89 staining).

**Figure 10.**
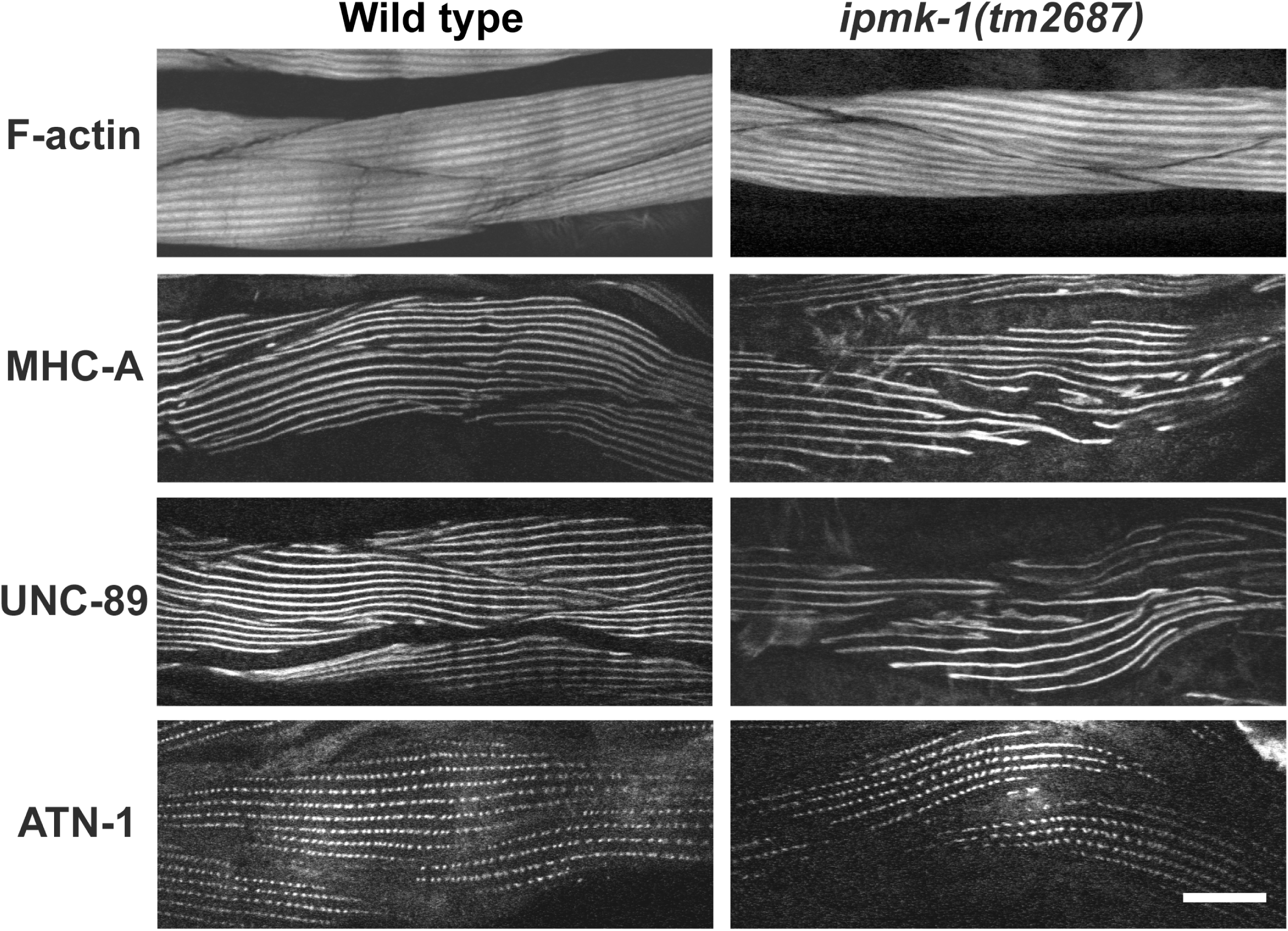
Mild disorganization of A-bands and M-lines in *ipmk-1(tm2687).* Confocal images of wild type and *ipmk-1(tm2687)* body wall muscle cells reacted with phalloidin (I-bands), and antibodies to MHC A (A-bands), UNC-89(M-lines) and ATN-1 (main portion of dense bodies). In *ipmk-1(tm2687),* although I-bands and dense bodies are not affected, there is mild disorganization of A-bands and M-lines, in that they are wavy and do not display the normal straight and parallel lines as seen in wild type. Scale bar, 10 µm.

We wondered if the MCB defect of *ipmk-1(tm2687)*, namely the large aggregates of IAC components, would affect muscle function. Thus, we performed quantitative motility assays, comparing wild type with *ipmk-1(tm2687),* outcrossed 6X to wild type. As shown in Figure 11B, *ipmk-1* is slower than wild type in both swimming and crawling, suggesting, that like *pix-1* pathway mutants (Moody et al., 2020, 2024), defects in MCB organization can reduce lateral transmission of forces required for optimal whole animal locomotion.

**Figure 11.**
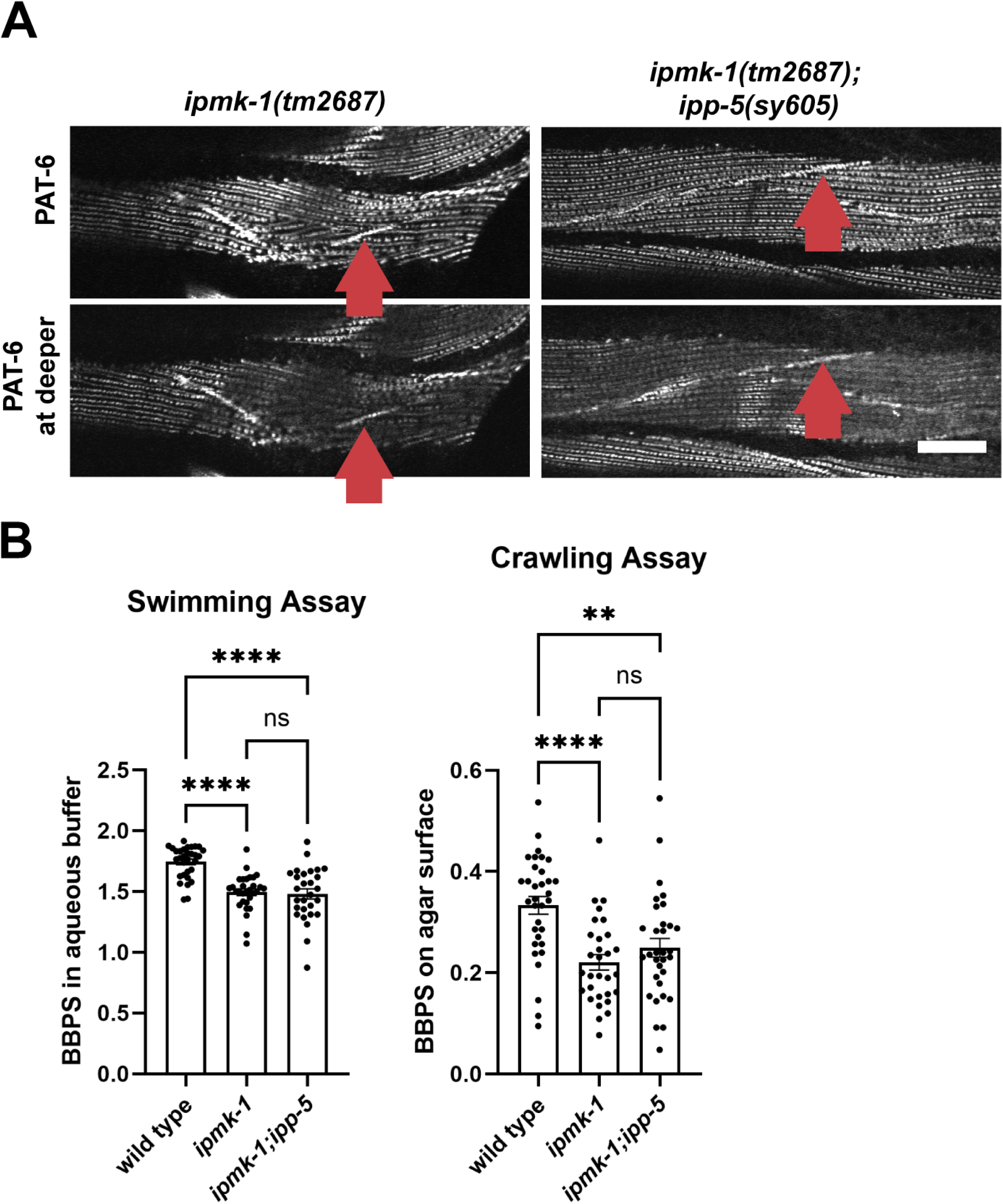
*ipp-5* fails to improve the MCB defect and locomotion defects of *ipmk-1*. A. anti-PAT-6 immunostaining of *ipmk-1(tm2687)* and *ipmk-1(tm2687); ipp-5(sy605)*. On top are shown images close to the outer muscle cell membrane; on the bottom are shown images deeper into the muscle cells. Note the clumping or the discontinuous distribution of PAT-6 at the MCB in either the single or double mutant (indicated by red arrows). Scale bar, 10 μm. B. Swimming and crawling assays of day 1 adults of the indicated genotypes. Note that in both swimming and crawling *ipmk-1* mutant animals are slower than wild type, and there is no improvement in swimming and crawling in *ipmk-1; ipp-5* double mutant animals. Statistical significance was tested using Brown-Forsythe and Welch ANOVA tests. ****: p<0.0001; **: p=0.0046; ns: not significant.

Yang et al. (2021) reported that *ipmk-1(tm2687)* has retarded post-embryonic growth, and a prolonged defecation cycle through a defect in Ca^2+^ signaling. We have confirmed the retarded growth but have not examined defecation. Interestingly, Yang et al. (2021) also reported that the growth and defecation defects can be rescued by loss of function of *ipp-5*. *ipp-5* encodes an inositol 5-phosphatase, that dephosphorylates IP3 to IP2 but does not convert PIP3 to PIP2. We created a double mutant between *ipmk-1(tm2687)*(outcrossed 6X to wild type) and a deletion allele of *ipp-5*, *ipp-5(sy605).* As shown in Figure 11A, *ipmk-1(tm2687); ipp-5(sy605)* animals have the same MCB defect as *ipmk-1(tm2687)* itself—abnormal clumping of PAT-6. In addition, these same *ipmk-1(tm2687); ipp-5(sy605)* double mutant animals are similarly defective in whole animal locomotion to *ipmk-1(tm2687)* itself, in both swimming and crawling (Figure 11B). Therefore, although *ipp-5* can rescue the growth (which we confirmed) and defecation defects, it cannot rescue the MCB structural defect and muscle function defect of *ipmk-1*. *ipp-5* is involved in regulating the level of IP3, a well-described regulator of Ca^2+^ homeostasis. Therefore, the lack of rescue of *ipmk-1* by *ipp-5*, suggests that the function of IPMK-1 at the MCB does not involve the Ca^2+^ role of IP3, but rather the cell membrane role of PIP3.

To determine the sub-cellular localization of IPMK-1, we raised rabbit polyclonal antibodies to it. As shown in Figure 12A, IPMK-1 is only 297 residues long, and by NCBI Conserved Domains, is predicted to have an IPK domain. We raised antibodies to the N-terminal most 119 residues of IPMK-1. As indicated in Figure 12B, these affinity-purified antibodies detect a protein of ∼34 kDa, the size predicted for IPMK-1, from wild type but not from two *ipmk-1* mutants, the original mutation found in the VC20386 MMP strain, *ipmk-1(gk214790)* which has a splice site mutation, and from *ipmk-1(tm2687)*, an intragenic deletion. Thus, our antibody is specific for IPMK-1, and both *ipmk-1* mutants are functionally protein nulls. We next used these antibodies to localize IPMK-1 in muscle. As shown in Figure 12C, in wild type body wall muscle IPMK-1 is found in discrete puncta which seem to be randomly distributed, not localizing near dense bodies or M-lines. Nevertheless, they appear in the same focal plane as the outer muscle cell membrane, similar to the location of PAT-6. In contrast, the *ipmk-1* deletion mutant shows only background staining with no clear or bright puncta. Images of these IPMK-1 puncta in wild type but not the *ipmk-1* null mutant can be seen more clearly in the enlarged views shown in Supplementary Figure 3.

**Figure 12.**
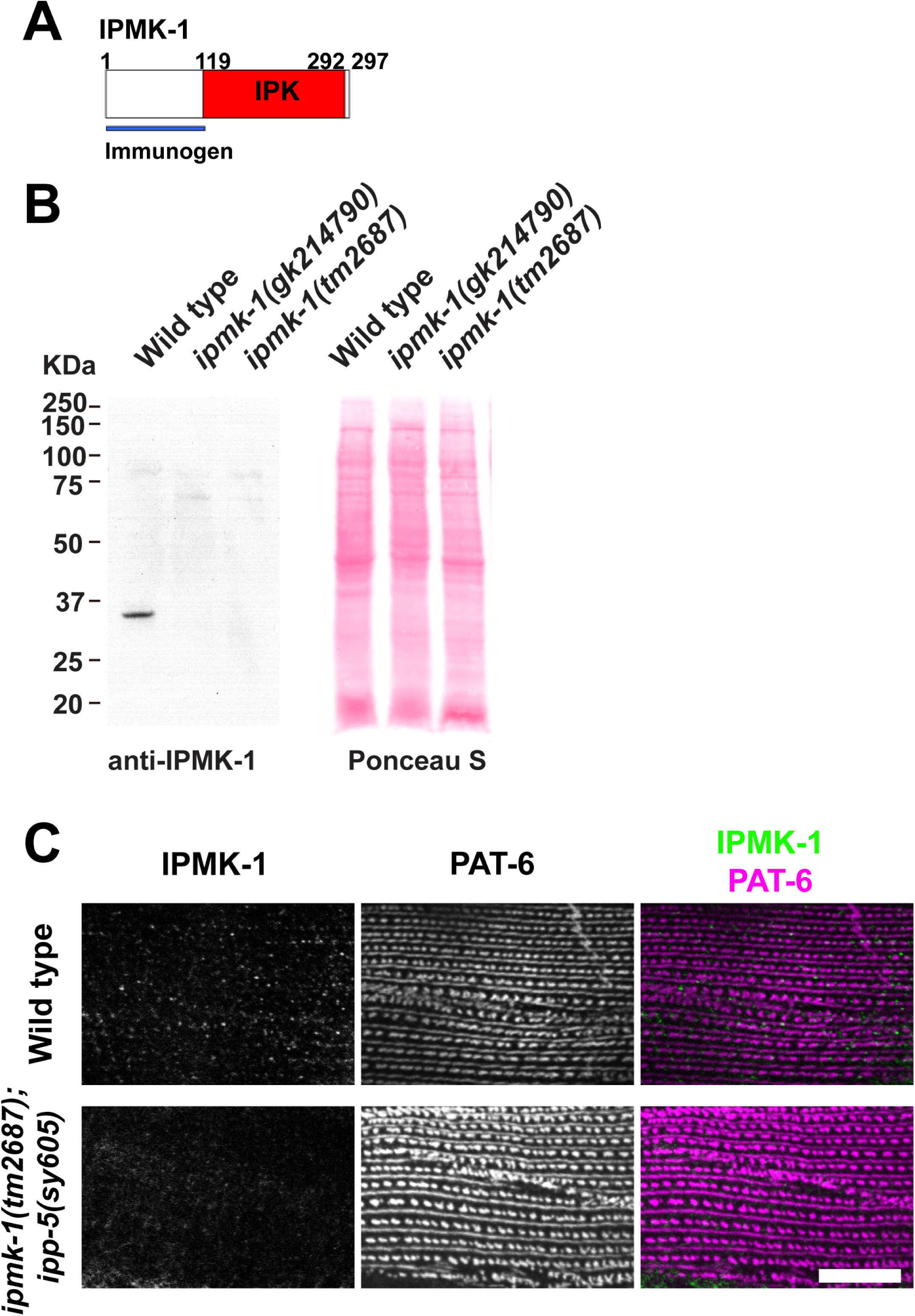
Antibodies to IPMK-1 localize IPMK-1 in body wall muscle cells to discrete puncta that are close to the muscle cell membrane but otherwise randomly distributed. A. Schematic of the IPMK-1 protein sequence, with the IPK domain predicted by NCBI Conserved Domains and location of immunogen. B. Western blot demonstrating detection of the expected 34 kDa IPMK-1 protein in wild type but not in splice site (*gk214790*) and deletion (*tm2687*) alleles of *ipmk-1*. The weak bands around 70-90 kDa are likely bacterial proteins. C. Co-immunostaining of wild type and *ipmk-1(tm2687); ipp-5(sy605)* with anti-IPMK-1 and anti-PAT-6. Note that in wild type IPMK-1 is found in discrete puncta in the same plane of focus as PAT-6, but otherwise randomly distributed. The signal from *ipmk-1; ipp-5* was much weaker. *ipp-5(sy605)* was used to overcome the growth defect of *ipmk-1(tm2687)* and thus provide larger muscle cells to image. Scale bar, 10 μm.

## Discussion

We began this study with the observation that the original MMP strain from which *pix-1* was identified (Moody et al. 2020), had a more severe muscle phenotype than the *pix-1* mutant that had been derived from it after several outcrosses. We then tested the hypothesis that lurking in the background of the original strain there was a genetic enhancer of the *pix-1* phenotype and that it was due to mutation in a single recessive gene. Our speculation was correct, and we were able to genetically map the enhancer to *ipmk-1*, which encodes inositol phosphate multikinase, an enzyme that converts PIP2 to PIP3, and IP3 to IP4, and IP4 to IP5.

PIX-1 is a RacGEF that promotes the exchange of GDP for GTP thereby activating Rac or Cdc42 and leading to activation of the protein kinase PAK-1. Loss of function mutations of *pix-1* and other members of the known *pix-1* pathway (e.g*. pak-1*, *ced-10* (Rac) and *rrc-1*) result in lack of accumulation of IAC components at the muscle cell boundary (MCB)(Moody et al. 2020, 2024). *ipmk-1; pix-1* double mutants show enhancement in that there is still lack of accumulation of IAC components (PAT-6, UNC-95, UNC-52) at the MCB but in addition, there is a large gap between adjacent muscle cells. Loss of function mutations in at least two other components of the pix pathway, *pak-1* and *rrc-1*, can also be enhanced by loss of function of *ipmk-1*. We would like to speculate that the gap is generated by the additive effects of lack of IAC assembly or stability at the MCB (the effect of mutations in the PIX pathway), and reduced attachment of M-lines and dense bodies to the ECM near the muscle cell boundaries due to altered PIP2/PIP3 ratios (the effect of the *ipmk-1* mutation). In support of this idea, our imaging of IAC components does show some mild disorganization of the regular row of dense bodies and the parallel alignment of M-lines near the MCB.

Loss of function of *ipmk-1* itself results in abnormal clumping of IAC components (PAT-6, UNC-112, UNC-95), rather than the normal uniform distribution of these components along the MCB arc. This clumping is likely due to either a deficiency in PIP3 or increased PIP2 in the muscle cell membrane. This idea is supported by the fact that mammalian orthologs of the IAC components UNC-112 (kindlin), TLN-1 (talin) and PAK-1 (PAK) have been demonstrated to bind to PIP3 and PIP2 (see below).

Yang et al. (2021) reported that *ipmk-1(tm2687)* has retarded postembryonic development and a prolonged defecation cycle. They provide evidence that this is due to a defect in Ca^2+^ signaling because it can be suppressed by providing extra Ca^2+^ in the growth media, or by a mutation in *ipp-5*, which specifies an inositol 5-phosphatase. IPP-5 converts IP3 to IP2, but not PIP3 to PIP2. Because our *ipmk-1; ipp-5* double mutant does not suppress the aggregation of PAT-6 at the MCB or the reduced whole worm locomotion (see below), we suggest that the function of IPMK-1 at the MCB does not involve Ca^2+^ signaling via IP3, but rather a deficiency of PIP3 or an abnormal ratio of PIP2 to PIP3 in the muscle cell membrane.

*ipmk-1* mutants display reduced whole animal locomotion likely due to reduced attachment of adjacent muscle cells to each other secondary to the non-uniform distribution of IAC components, and consistent with a role of lateral force transmission at MCBs for optimal whole animal locomotion. This explanation was previously suggested for our observation that *pix-1* and *rrc-1* mutants display reduced whole animal locomotion (Moody et al. 2020, 2024). *ipmk-1* also results in mild disorganization of the other IAC structures, M-lines and dense bodies, and also mild disorganization are sarcomeric A-bands and M-lines. These later defects may also contribute to the observed reduction in whole animal locomotion.

An antibody to IPMK-1 detects the protein by western blot from wild type but not from two *ipmk-1* mutant alleles and localizes to puncta near the outer muscle cell membrane but not in an organized pattern, and not coincident with MCBs, M-lines or dense bodies. This localization pattern is consistent with the idea that IPMK-1 generates PIP3 in the cytosol near the muscle cell membrane which then inserts into it.

There is an extensive body of literature demonstrating the role of phosphoinositides in IAC assembly in mammalian non-muscle cells. Protein interactions forming IACs, and their association with the cell membrane display features of the in-vogue concept of membrane-associated biomolecular condensation facilitated by liquid-liquid phase separation (Case et al. 2019; Hsu et al. 2023; Hsu et al. 2025). IAC assembly itself is influenced by the composition of the cell membrane lipids (Legate et al. 2011; Tachibana et al. 2023). Although phosphoinositides comprise a minor fraction of membrane lipids, (PIP2 at 1-3%, and PIP3 at 0.01—0.05% (Insall et al. 2001)), PIP2 and PIP3 influence membrane dynamics and integrin activation (DiPaolo et al., 2006). Important IAC components, kindlin (UNC-112) and talin, involved in integrin activation and thus IAC formation, bind to these PIPs. Kindlin binds to PIP2 and PIP3 through its PH domain, and at least for kindlin-3, with somewhat higher affinity for PIP3, and binding is facilitated by clustering of PIP3 (Liu et al. 2011; Ni et al. 2017). PIP3-mediated binding of kindlin-2 was shown to be critical for integrin activation because wild type kindlin-2 enhanced talin-mediated activation of integrin, but PIP3 binding-defective kindlin 2 failed to enhance talin mediated integrin activation in CHO cells (Liu et al. 2011). Talin binds to PIP2 through its FERM domain (Chinthalapudi et al. 2018). In fact, PIP2 has been reported to recruit kindlin and talin to the cell membrane and stimulate localized phase separation of newly forming IACs (Hsu et al., 2023).

Interestingly, the FERM domain of talin binds to and thereby stimulates the enzymatic activity of phosphatidyl-inositol phosphate kinase type 1γ (PtdInsPKIγ), which generates PIP2. Moreover, PtdInsPKIγ is concentrated at focal adhesions (IACs) of NIH 3T3 cells and at the IACs of neuronal synapses. Overexpression of PtdInsPKIγ disrupts IACs at least in NIH 3T3 cells (Di Paolo et al. 2002). The talin binding site of PtdInsPK1γ localizes the enzyme to focal adhesions (IACs)(Di Paolo et al. 2002; Ling et al. 2002). Legate et al. (2011) cleverly deleted an exon that encodes the talin-binding focal adhesion targeting sequence from mouse embryonic stem cells and used it to establish a mouse strain, and then studied embryonic fibroblast and kidney fibroblasts from these mice, but mostly studied the kidney fibroblasts. The recruitment of talin and vinculin but not kindlin 2 were impaired in the focal adhesion targetless PtdInsPK1γ cells. In addition, the focal adhesion targetless PtdInsPK1γ cells showed initially a reduced rate of cell attachment via integrin-fibronectin, and this was corrected within minutes but integrin-actin force coupling remained defective.

The generation of PIP3 by phosphoinositide-3-OH kinase (Pi(3)K) also influences integrin linked kinase (ILK; PAT-4 in worms). Delcommenne et al. (1998) reported that ILK kinase activity is stimulated by PIP3, and that in tissue culture cells, insulin stimulated and fibronectin attachment stimulated ILK kinase activity is dependent on generation of PIP3 since these activities are blunted upon chemical inactivation of Pi(3)K. Moreover, transfection of constitutively activated Pi(3)K results in higher ILK activity.

Finally, the IAC component vinculin (DEB-1 in worms) binds to PIP2 via basic residues in its C-terminal tail domain. Although PIP2 binding is not required for localization of vinculin to IACs (focal adhesions), it is required for vinculin activation and turnover of IACs in mouse fibroblasts (Thompson et al. 2017).

There is precedence for a connection between PIP2 and the PIX pathway. The known final component of the PIX pathway, PAK (in this case human PAK1), through a positively charged region upstream of its Rac/Cdc42 binding region, binds to PIP2 of the cell membrane and further enhances the kinase activity of already GTP-Rac or GTP-Cdc42 bound PAK1 (Strochlic et al. 2010).

So far, all these interesting studies in the literature have been conducted on tissue culture cells. Our results demonstrate the importance of PIP2/PIP3 levels for proper IAC formation in vivo, and in particular, in muscle, by characterization of a loss of function mutant in a PIP3-generating enzyme, IPMK-1. Moreover, the PIP binding IAC proteins, kindlin and talin, have orthologs in worms, UNC-112 (Rogalski et al. 2000) and TLN-1 (Moulder et al. 1996; Cram et al. 2003), respectively. In addition, the conserved PIX scaffolding protein, GIT (GIT-1 in worms) has an ARFGAP domain that is known to bind to PIP3 to localize it to the cell membrane (Czech 2000). Finally, we have speculated that the PIX pathway member, RacGAP RRC-1, is membrane-associated, and we have recently found that RRC-1 contains a PX domain which is known to bind to PIP3 (Sagadiev et al., in preparation).

## Data availability

The authors declare that the data supporting the results of this study are available within the paper or within the supplementary material, and additional details are available upon request.

## Acknowledgements

We thank Robert Barstead (Oklahoma Medical Research Foundation) for the cDNA library RB2, Andrew Fire (Stanford University) for the *myo-3* promoter plasmid pPD95.86, and Pamela Hoppe (Western Michigan University) for monoclonals MH35 and MH42. Most of the nematode strains used in this work were provided by the Caenorhabditis Genetics Center, which is funded by the National Institutes of Health Office of Research Infrastructure Programs (P40OD010440). Some nematode strains were obtained from Shohei Mitani at the National BioResource Project (*C. elegans*) at Tokyo Women’s Medical University School of Medicine.

## Study funding

The authors gratefully acknowledge support from the National Institutes of Health, grant R01HL160693 to G.M.B.

## Conflict of interest

The authors declare no conflicts of interest.

---

**Supplementary Figure 1. The enhancer mutation is not on chromosome III.** Since we used a GFP insertion in chromosome III to follow our crosses, we next crossed en/en; *pix-1/pix-1* X 3xC with a strain in which GFP had been inserted in chromosome IV, thus eliminating mutations on chromosome III. Upon immunostaining, the resulting strain still showed the “big gap” phenotype, suggesting that the enhancer mutation does not reside on chromosome III but that the enhancer mutation does reside on the left arm of chromosome IV. Scale bar, 10 µm.

**Supplementary Figure 2. Double mutant analysis shows that 4 genes, when mutant, do not enhance the phenotype of *pix-1*.** The indicated double mutants were made with *pix-1*. These 4 genes (*alp-1*, C27B7.7, K08E7.6, and K07F5.8) were candidates for the enhancer, in addition to *ipmk-1*, as shown in Figure 5. Shown are representative examples of immunostaining of body wall muscle with anti-PAT-6. Note that none of the double mutants show the large gap phenotype.

**Supplementary Figure 3. Enlargement of Figure 12C anti-IPMK-1 immunostaining.** Portions of 2 body wall muscle cells from either wild type or *ipmk-1(tm2687); ipp-5(sy605)* immunostained with antibodies to IPMK-1 (green) and anti-PAT-6 (magenta). *ipp-5(sy605)* was used to overcome the growth defect of *ipmk-1(tm2687)* and thus provide larger muscle cells to image. Note the puncta of IPMK-1 in wild type but not in *ipmk-1(tm2687); ipp-5(sy605)*. Scale bar, 10 μm.

**Supplementary Table 1. PCR primers used to amplify regions that contain SNPs in VC20386 as compared to wild type.**

**Supplementary Table 2. PCR primers used to narrow down the region on IV that contains the enhancer mutation.**

**Supplementary Table 3. Information about 11 genes that contain mutations in en/en; *pix-1/pix-1* 3X OC strain.**

